# Integrated Short-TE and Hadamard-edited Multi-Sequence (ISTHMUS) for Advanced MRS

**DOI:** 10.1101/2024.02.15.580516

**Authors:** Steve C.N. Hui, Saipavitra Murali-Manohar, Helge J. Zöllner, Kathleen E. Hupfeld, Christopher W. Davies-Jenkins, Aaron T. Gudmundson, Yulu Song, Vivek Yedavalli, Jessica L Wisnowski, Borjan Gagoski, Georg Oeltzschner, Richard A.E. Edden

## Abstract

**Background:** To examine data quality and reproducibility using ISTHMUS, which has been implemented as the standardized MR spectroscopy sequence for the multi-site Healthy Brain and Child Development (HBCD) study.

**Methods:** ISTHMUS is the consecutive acquisition of short-TE PRESS (32 transients) and long-TE HERCULES (224 transients) data with dual-TE water reference scans. Voxels were positioned in the centrum semiovale, dorsal anterior cingulate cortex, posterior cingulate cortex and bilateral thalamus regions. After acquisition, ISTHMUS data were separated into the PRESS and HERCULES portions for analysis and modeled separately using Osprey. In vivo experiments were performed in 10 healthy volunteers (6 female; 29.5±6.6 years). Each volunteer underwent two scans on the same day. Differences in metabolite measurements were examined. T_2_ correction based on the dual-TE water integrals were compared with: 1) T_2_ correction based the default white matter and gray matter T_2_ reference values in Osprey; 2) shorter WM and GM T_2_ values from recent literature; and 3) reduced CSF fractions.

**Results:** No significant difference in linewidth was observed between PRESS and HERCULES. Bilateral thalamus spectra had produced significantly higher (p<0.001) linewidth compared to the other three regions. Linewidth measurements were similar between scans, with scan-to-scan differences under 1 Hz for most subjects. Paired t-tests indicated a significant difference only in PRESS NAAG between the two thalamus scans (p=0.002). T_2_ correction based on shorter T_2_ values showed better agreement to the dual-TE water integral ratio.

**Conclusions:** ISTHMUS facilitated and standardized acquisition and post-processing and reduced operator workload to eliminate potential human error.

**Highlights:**

- ISTHMUS has been implemented into the HBCD study protocol.
- It acquires both short-TE and Hadamard-edited transients.
- ISTHMUS reduces operator workload.
- ISTHMUS potentially allows improved T2 relaxation correction

## 1 INTRODUCTION

In vivo MRS allows for non-invasive quantification of brain metabolites but is limited by spectral resolution due to broad in vivo linewidths (in the order of 10 Hz) and peak splitting due to J-couplings. The limited dispersion of signals along the chemical shift dimension leads to spectral overlap particularly for metabolites with low concentration. Edited MRS improves spectral resolution by reducing the number of signals with frequency-specific manipulation of known couplings in target metabolites (such as the inhibitory neurotransmitter γ-aminobutyric acid GABA and the antioxidant glutathione GSH). It has recently been accelerated by Hadamard-encoded editing including the HERMES and HERCULES experiments (Chan et al., 2016; Chan et al., 2017; Oeltzschner et al., 2019; Saleh et al., 2016). The HERCULES sequence measures multiple low concentration metabolites simultaneously. It is a four-step Hadamard-encoded editing scheme for simultaneous editing and linear combination modeling (LCM) of low-concentration metabolites including ascorbate (Asc), aspartate (Asp), γ-aminobutyric acid (GABA), glutathione (GSH), lactate (Lac), N-acetylaspartate (NAA) and N-acetylaspartylglutamate (NAAG), reported as tNAA (NAA+NAAG) and phosphorylethanolamine (PE), as well as a couple of high-concentration metabolites including Glutamine (Gln) and Glutamate (Glu), reported as Glx (Gln + Glu), myo-inositol (mI), total Creatine (tCr = Cr + PCr) and total Choline (tCho = GPC + PCh). 20-ms editing pulses are applied at: (A) 4.58 and 1.9 ppm; (B) 4.18 and 1.9 ppm; (C) 4.58 ppm; and (D) 4.18 ppm. More details are available in (Oeltzschner et al., 2019). A major drawback of edited MRS is that longer TEs are required for optimized editing of the target metabolites. This results in T_2_ decay of metabolite signals, which in turn results in reduced metabolite signalLJtoLJnoise ratio (SNR) and potential misinterpretation of relaxation changes as concentration changes.

In contrast, unedited MRS for high-concentration metabolite measurements can be achieved using a short TE typically between 30 and 35 ms using PRESS localization (Bottomley, 1987). PRESS is a single-shot localization sequence consisted of one slice-selective 90° excitation pulse and two slice-selective 180° refocusing pulses, each applied concurrently with one of three orthogonal gradients (x, y and z) (Bottomley, 1987). The two refocusing pulses yield a spin echo that fully utilizes the available M_z_ magnetization to produce a full-intensity localized signal (Hamilton et al., 2009; Zhong and Ernst, 2004) for an SNR-efficient acquisition. Spectra acquired at short TE are characterized by higher SNR due to lesser T_2_ decay and more pronounced metabolite resonances due to lesser J-evolution. However, underlying mobile macromolecules (MM) and lipid resonances distort the metabolite spectra at short TEs (Cudalbu et al., 2012). This artifact can be minimized by advanced post-processing with successful baseline correction and inclusion of a cohort-mean MM basis function during curve fitting (Zollner et al., 2023b). At short TE, Glx (glutamine + glutamate) and GABA show multiple resonance peaks between 2.05 and 2.45 ppm partially overlapped by the NAA multiplet making it difficult to resolve the peaks from one another. The above-mentioned editing techniques are useful to probe these metabolites.

T_2_-weighting of the MRS spectrum increases with TE. Metabolites (e.g. mI, Glx and lipid) and MM with short T_2_ relaxation times are easier to detect in short-TE spectra and these signals decay with longer TE (Cudalbu et al., 2021; Mlynarik et al., 2001). A short TE also allows acquisition of water references and metabolites signals with lesser T_2_ weighting. This is particularly important for T_2_-corrected metabolites measurements in LCM as details of metabolites’ T_2_s are limited, especially for young children. In current practice, age-group-specific relaxation times are not considered in most fitting tools when T_2_ correction is performed. Metabolite measurements on a shorter TE are therefore more accurate due to a reduced dependence on assumed metabolite T_2_ values [ref Mullins paper].

In vivo brain metabolite T_2_ relaxation times have been reported from a few studies. Most early studies reported the relaxation times of prominent singlet signals such as tNAA, tCr and tCho at 1.5T and 3.0T (Barker et al., 2001; Brief et al., 2005; Mlynarik et al., 2001; Traber et al., 2004; Zaaraoui et al., 2007). Two other studies performed the measurements of low-concentration metabolites with complex coupled spin systems including Asp, GABA, Glx, GSH, mI etc. (Ganji et al., 2012; Wyss et al., 2018) and at different field strengths including 4.0T, 7.0T and 9.4T (Marjanska et al., 2012; Michaeli et al., 2002; Murali-Manohar et al., 2020). Most metabolite T_2_ relaxation values are reported based on adults. Comparative studies have reported inconsistent results for the change in metabolite T_2_ relaxation for NAA, tCr and tCho between young and elderly individuals (Brooks et al., 2001; Marjanska et al., 2013). The discrepancy may result from data collected at different field strengths and from different brain regions. Pediatric metabolite T_2_ has been rarely studied - one study included a small group of adolescents which reports the T_2_ of NAA, Cho and Cr (Kirov et al., 2008). This gap in knowledge likely arises from the longer acquisition duration of experiments to measure T_2_ relaxation times, which are difficult to perform in pediatric populations. Significantly different T_2_ values between singlet and multiplet resonance groups can be a confound for linear combination modeling of medium-TE data (Soher et al., 2005).

The purpose of this study was to present the Integrated Short-TE and Hadamard-edited Multi-Sequence (ISTHMUS) which combines a short-TE unedited PRESS (high SNR, low T_2_-weighting, low spectral resolution for low concentration metabolites of interest) and a long-TE edited HERCULES (lower SNR, higher T_2_-weighting, prioritizing spectral resolution of targeted low concentration metabolites of interest) into a single scan, delivering a complementary set of metabolic information in a single acquisition. It has been implemented into the full protocol for the HEALthy Brain and Child Development (HBCD) study (Jordan et al., 2020), a longitudinal study that follows children’s developing brains from birth through early childhood. A feature of this integrated sequence is that it combines the united and edited sequences with dual-TE water reference scans into one single acquisition. As there are benefits and costs of short- and long-TE acquisitions, data acquired using both TEs in a single acquisition may mitigate the various drawbacks of each. The ISTHMUS acquisition also offers the potential to investigate T_2_ relaxation for water and metabolites by combining the short- and long-TE acquisitions. With very limited data and knowledge on metabolite T_2_ relaxation times especially on neonates, a multi-TE sequence is helpful to gather more information on the T_2_ decay and hence is more robust to T_2_ changes. In this study, we present data and quantification of metabolite concentrations acquired using ISTHMUS, test the quality and measure the consistency of them, and describe the benefits of the sequence.

## 2 METHODS

ISTHMUS is the consecutive acquisition of short-TE PRESS and long-TE HERCULES data in the same volume of interest. Additionally, dual-TE water reference scans are collected during the ISTHMUS acquisition as shown in Figure 1. All analyses and results reported in this study are based on the implementation of ISTHMUS on Philips 3T MRI scanners. For clarity of communication, ISTHMUS data that are short-TE unedited PRESS will be referred to as ‘PRESS’ data and the long-TE edited HERCULES will be referred to as ‘HERCULES’ data in this manuscript. In ISTHMUS, based on the implementation shown in Figure 1b, 32 short-TE PRESS transients are acquired in a block at the start (TE 35 ms, FA 90 °, shown in green), followed by 224 transients of HERCULES (TE 80 ms, shown in orange). 4 transients of short-TE (35 ms) water reference signal and 4 transients of long-TE (80 ms) water reference signal (both shown in blue) are interleaved throughout the acquisition. Real-time frequency correction for field drift is performed based on the water frequency of the reference scans from the localized voxel, which are acquired every 32 transients (Edden et al., 2016). Additional acquisition parameters include: 3T field strength; TR 2000 ms; second-order auto shim, PRESS excitation/refocusing bandwidth 2200/1350 Hz; 2048 datapoints sampled at 2 kHz. Full scan parameters are reported in the Supplementary Material 1 according to the minimum reporting standards for in vivo magnetic resonance spectroscopy (MRSinMRS) checklist (Lin et al., 2021).

**Figure 1a.**
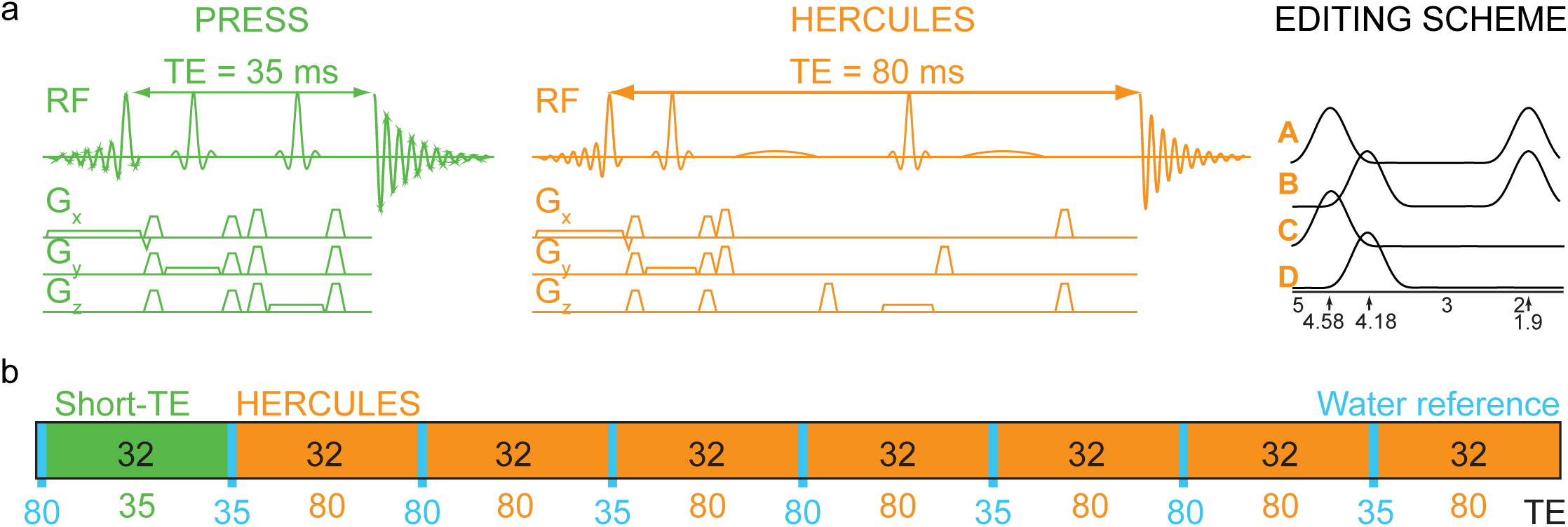
ISTHMUS combines a 35 ms short-TE unedited scheme (green) and a 80 ms long-TE edited HERCULES scheme (orange) using PRESS localization. Four 20-ms editing pulses (black) targeting 1.9 ppm, 4.18 ppm and 4.56 ppm are applied for Hadamard combinations. 1b. ISTHMUS multi-sequence schedule starting with 32 transients of short-TE acquisition (green) followed by 224 transients of long-TE edited HERCULES scheme (orange). 4 long- and 4 short-TE interleaved water reference scans (blue) are acquired evenly throughout the sequence.

### Phantom Experiments

ISTHMUS experiments were performed using a 1000 mL cylindrical phosphate-buffered saline phantom containing Cho (3mM), Cr (10mM), GABA (2mM), L-glutamic acid (10mM), L-glutamine (3mM), Lac (5mM), myo-inositol (7.5 mM), NAA (12.5 mM) and sodium azide (0.02%) at room temperature on a Philips 3T MRI scanner (Ingenia CX, Philips Healthcare, The Netherlands) using a 32-channel phased-array head coil. Scan parameters were the same as introduced in the previous section in a 3×3×3 cm^3^ voxel positioned in the middle of the phantom at iso-center. Philips ‘excitation’ water suppression was applied after optimization. The duration of the scan was approximately 9 minutes.

### In Vivo Experiments

ISTHMUS experiments were performed in 10 healthy volunteers (6 females and 4 males; aged 23-46; mean 29.5 ± 6.6 years) with local IRB approval in 4 brain regions using the same Philips 3T scanner and head coil as described in the phantom experiments with CHESS water suppression. T_1_-weighted MPRAGE (TR/TE/ 6.9/3.2 ms; FA 8°; 1 mm^3^ isotropic resolution) MRI was acquired for voxel positioning and tissue segmentation. Voxels were localized in the predominantly white matter centrum semiovale (CSO), the predominantly gray matter posterior cingulate cortex (PCC) both in the size of 30×26×26 mm^3^, dorsal anterior cingulate cortex (dACC) 30×30×30 mm^3^ and bilateral thalamus 25×36×25 mm^3^ as shown in Figure 2. Participants completed two consecutive 39-minute scan protocols, separated by a short (∼10 minutes) removal from the scanner, for examining test-retest reproducibility. In the second scan, voxels were positioned in the same brain region by visual inspection, as close as possible to the first scan. Exclusion criteria included contraindications for MRI and a history of neurological and psychiatric illness.

**Figure 2.**
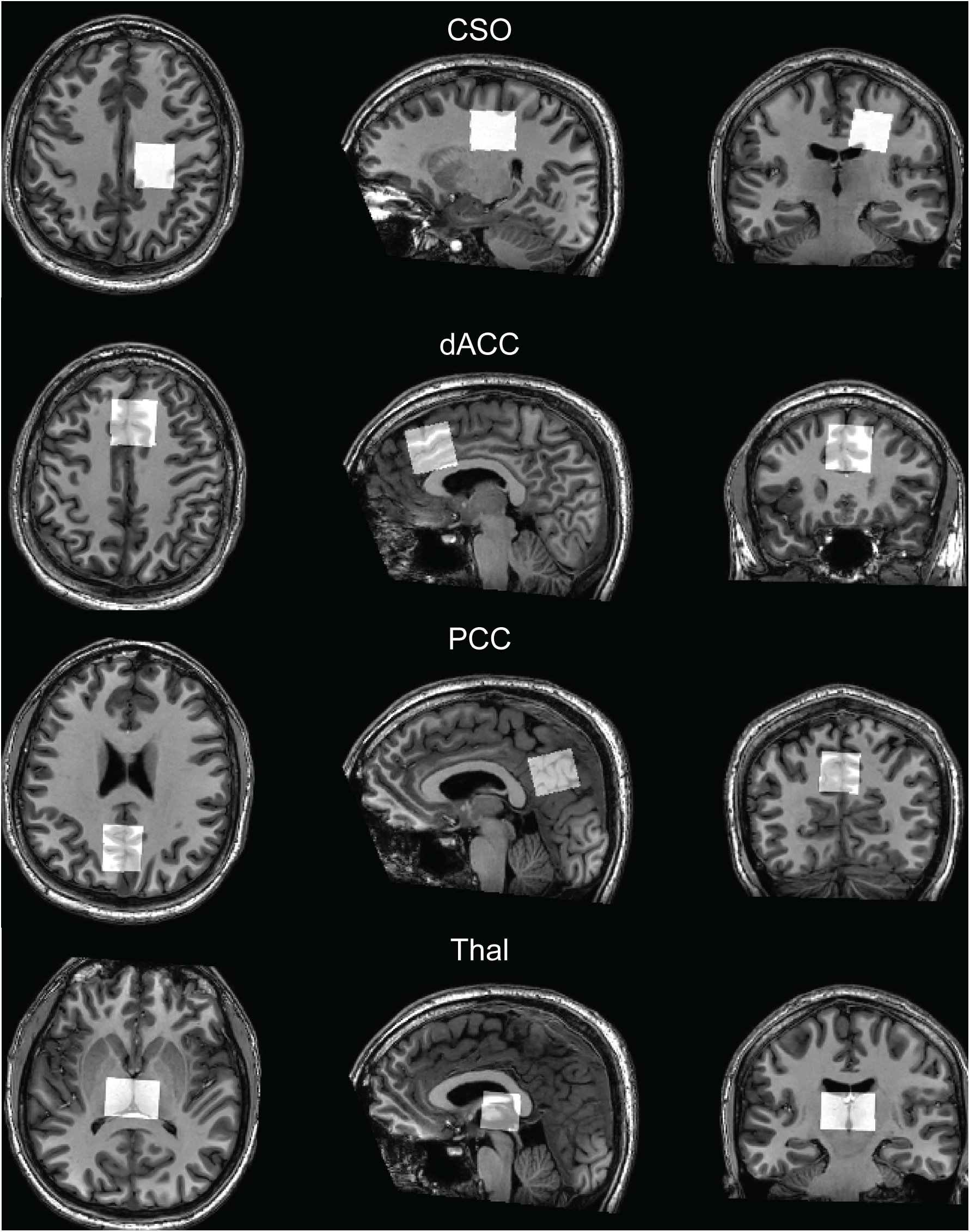
Voxel locations of CSO (30×26×26 mm^3^), dACC (30×30×30 mm^3^), PCC (30×26×26 mm^3^) and bilateral thalamus (25×36×25 mm^3^).

### Quantification

After acquisition, ISTHMUS data were separated into PRESS and HERCULES data using spec2nii (Clarke et al., 2022) followed by processing and linear combination modeling (LCM) using Osprey (5). LCM of the PRESS and HERCULES data, including each Hadamard combination spectrum (i.e. sum, difference-1 and difference-2) was performed separately in Osprey (i.e. no multi-spectrum modeling was performed). The sequence-specific basis sets for PRESS and HERCULES were generated from a fully localized 2D density-matrix simulation (101×101 spatial resolution) using real vendor pulse waveforms and sequence timings (Hui et al., 2022b; Simpson et al., 2017). Simulations were accelerated by the one-dimensional projection method (Zhang et al., 2017) and the use of coherence pathways filters (Bodenhausen et al., 2011). More details are available in (Hui et al., 2022b). The metabolite basis functions included ascorbate (Asc), aspartate (Asp), creatine (Cr), negative creatine methylene (-CrCH_2_), gamma-aminobutyric acid (GABA), glycerophosphocholine (GPC), glutamine (Gln), glutamate (Glu), glutathione (GSH), lactate (Lac), myo-inositol (mI), N-acetylaspartate (NAA), N-acetylaspartylglutamate (NAAG), phosphocholine (PCh), phosphocreatine (PCr), phosphoethanolamine (PE), scyllo-inositol (sI), and taurine (Tau). The selection of metabolites to be included was based on the neurochemical profiles observed in the healthy adult brain to avoid any quantification bias (Tkac et al., 2009). Cohort-mean measured mobile macromolecule (MM) basis functions were used (Zollner et al., 2022), derived as described previously (Hui et al., 2022a). The simulated metabolite and measured MM basis functions were incorporated into the Osprey software for modeling. Metabolite measurements were quantified relative to the water reference signal and tissue-corrected for tissue-specific water visibility and relaxation times based on literature values (Wansapura et al., 1999), with primary outcome measures based upon correction at each echo time separately without integration of multi-TE data (see also the following section). Osprey-integrated SPM12 (Friston et al., 1994) was used to perform brain tissue segmentation to yield relative tissue volume fractions of gray matter (GM), white matter (WM), and cerebrospinal fluid (CSF). Metabolite concentrations were calculated in Osprey, using the tissue corrections reported in (Gasparovic et al., 2006).

Visual inspection was performed to evaluate artifacts and contamination to ensure data quality according to consensus recommendations (Wilson et al., 2019). Datapoints that had a concentration measurement of 0 were excluded, interpreted as modeling failure. The water-reference spectra were modeled with a simulated water basis function in the frequency domain with a 6-parameter model (amplitude, zero-and first-order phase, Gaussian and Lorentzian line broadening, and frequency shift). For phantom data, reduced basis functions included Cr, GABA, Gln, Glu, Lac, mI, NAA, PCh and PCr were used for modeling; no lipid and MM basis functions were included.

### Investigation of water T_2_ relaxation correction

The collection of dual-TE water reference scans allows us to validate the T_2_ correction performed in Osprey. This illustrates a potential benefit of ISTHMUS acquiring metabolite and water data at multiple echo times. Dual-TE water reference integral ratios *R*_integral_ were calculated using Equation 1 and compared with the ratio of Osprey T_2_ correction factors calculated using Equation 2. The water reference integral ratios were calculated by:

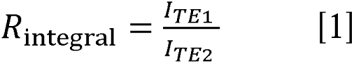

where *I_TE_*_1_ denotes the integral of the short-TE water reference signal and *I_TE2_*denotes the integral of the long-TE water reference signal.

The ratio *R*_segmentation_ between the Osprey T_2_ correction factors at short- and long-TE (i.e. 35 ms and 80 ms) was calculated from the segmented voxel tissue fractions and reference T_2_ values of GM, WM and CSF from the literature (Piechnik et al., 2009; Wansapura et al., 1999) thus:

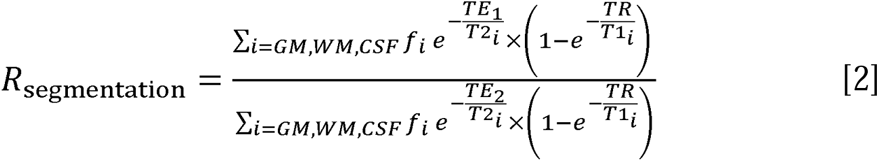

where *f_i_* denotes the molal signal fraction of each tissue compartment, T1_i_ and T2_i_ are literature-based relaxation time constants for each tissue compartment where *i* is GM, WM or CSF; TE_1_ and TE_2_ denote the short and long TEs, respectively. TR is the repetition time of 2000 ms. *R*_integral_, the water reference signal ratio was then plotted against *R*_segmentation_, the Osprey T_2_ correction-factor ratio in order to investigate the validity of the T_2_ relaxation correction that was applied for quantification above. Ideally, the two ratios should be equal, and yield a perfect correlation of slope one and intercept zero.

Alternative values of *R*_segmentation_ were also calculated using shorter T_2_ values for WM and GM from a more recent publication (Zi et al., 2020) to investigate the impact of errors in the reference T_2_ relaxation times. Lastly, values of *R*_segmentation_ were calculated again, with default Osprey T_2_ values and reducing the segmented voxel fraction of CSF by 50%, to investigate the impact of CSF segmentation errors.

Finally, metabolite concentrations were re-calculated using *R*_integral_ to infer a single water T_2_ for the voxel. This analysis is exploratory, and is not included in the primary outcome variables. The average fractional change in metabolite concentration with this approach was calculated, and the correlation coefficient was calculated for each metabolite between these concentrations and the primary outcome variables.

### Statistical Analysis

All statistical analyses were performed using R v4.0.2 in RStudio v1.2.5019 (RStudioTeam, 2020). Data quality metrics including SNR and linewidth of the 3-ppm tCr signal were calculated for the processed metabolite spectra from PRESS and HERCULES in CSO, dACC, PCC and bilateral thalamus. SNR is defined as the ratio of the tCr singlet peak height and the detrended standard deviation of the frequency-domain spectrum between −2 and 0 ppm. Linewidth was defined as the average of the peak-measured FWHM and the FWHM of a Lorentzian model of the tCr singlet. One-way ANOVA and the Tukey’s multiple comparisons correction were used to compare linewidths between PRESS and HERCULES datasets. Tissue-corrected metabolite concentrations were calculated for Asc, Asp, GABA, GSH, Lac, NAA, NAAG, tCho, tCr, Gln, Glu and mI. Coefficients of variation (CV) were calculated for each metabolite to compare the between-subject variation of metabolite measurements (i.e. CVs from the total number of measurements from 10 participants in 20 scans). Data collected from the first and second session of the scan were compared using paired t-test and visualized using a Bland-Altman plot to evaluate reproducibility of ISTHMUS. Correlation coefficients between the water reference ratios and the Osprey correction factors and their respective CVs were measured. P-values less than 0.05 were considered statistically significant. Adjusted p-values less than 0.004 (0.05/12) were considered statistically significant when repeated measurements were performed for targeted metabolites (Asc, Asp, GABA, GSH, Gln, Glu, mI, Lac, NAA, NAAG, tCho, tCr).

## 3 RESULTS

Phantom experiments showed successful data acquisition using ISTHMUS. The dataset was separated into PRESS and HERCULES data as shown in Figure 3. SNR and linewidth of tCr were 96.1 and 3.2 Hz for PRESS and 248.3 and 3.5 Hz for HERCULES. Metabolite concentration measurements in the phantom from ISTHMUS are shown in Table 1. NAA, PCh and tCr were comparable between the PRESS and HERCULES sum spectrum. GABA and Lac were clearly visible as shown in Figure 3c and 3d. Since the purpose of the phantom scan was to demonstrate the functionality of ISTHMUS, no quantifications and statistical analysis were performed for this part.

**Figure 3.**
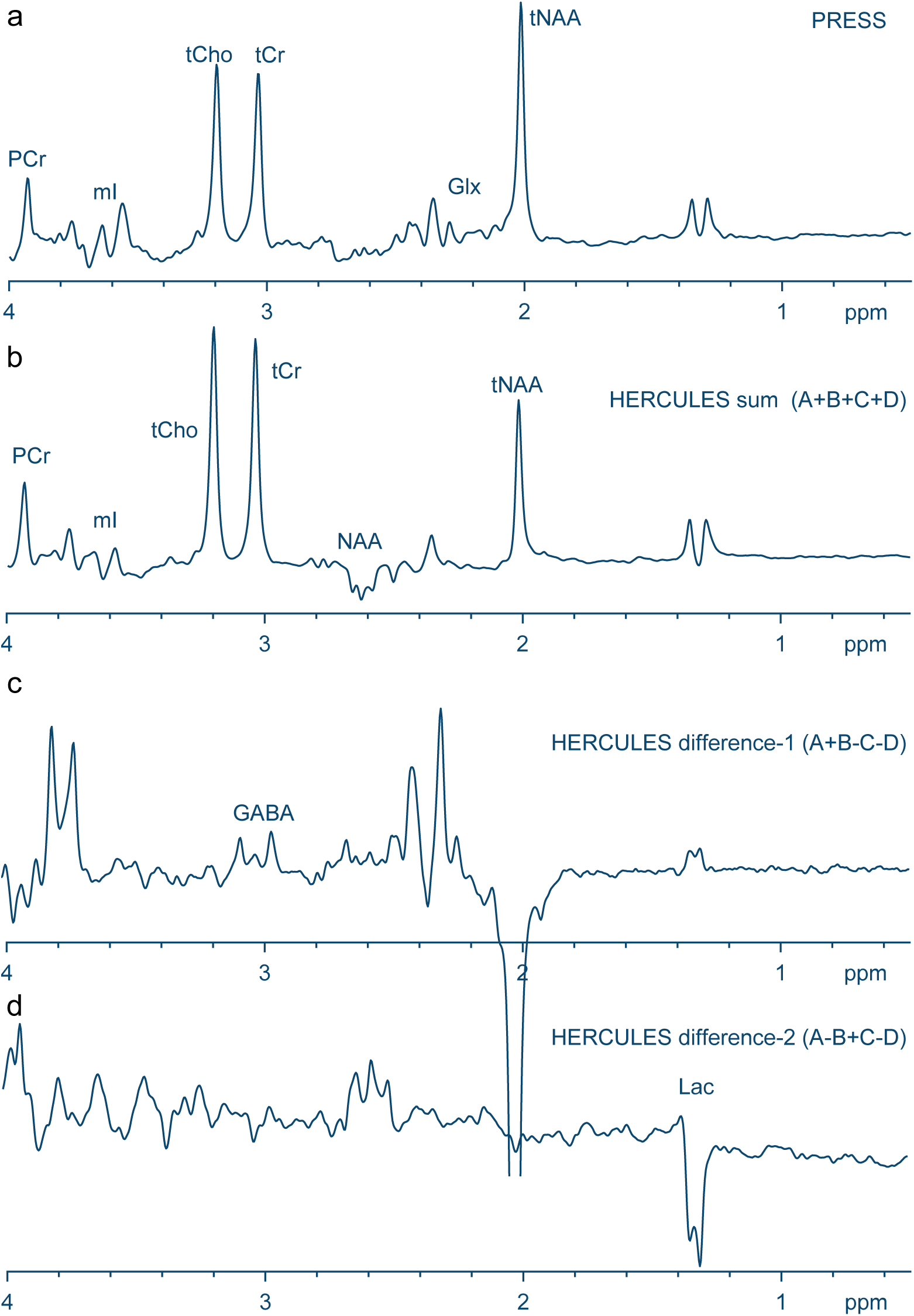
Modeling of the phantom data. 3a) Short-TE unedited PRESS and 3b) long-TE HERCULES sum clearly show conventional unedited metabolites. 3c) Hadamard combinations spectra for modeling GABA, and 3d) Hadamard combinations spectra for modeling Lac. The phosphate-buffered saline phantom contains Cho, Cr, GABA, Glu, Gln, Lac, mI and NAA. Asc, Asp, GSH and NAAG were not included.

**Table 1.** Quantitative water-scaled metabolite concentration measurements from the phantom data acquired using ISTHMUS. The data were separated into the PRESS (32 transients) and HERCULES (224 transients) compartments and modeled separately using Osprey. Gln and Glu were quantified separately and reported as Glx.

|  | Cr | GABA | Gln | Glu | mI | Lac | NAA | PCh | PCr | Glx | tCr |
| --- | --- | --- | --- | --- | --- | --- | --- | --- | --- | --- | --- |
| PRESS | 2.37 | 3.56 | 1.60 | 10.67 | 5.63 | 4.50 | 9.17 | 3.46 | 4.85 | 12.28 | 7.21 |
| HERCULES sum | 0.00 | 0.00 | 1.75 | 6.58 | 1.45 | 2.79 | 8.57 | 2.39 | 6.27 | 8.33 | 6.27 |
| HERCULES difference-1 | 0.84 | 0.68 | 1.77 | 8.07 | 1.31 | 6.18 | 8.08 | 0.00 | 1.10 | 9.84 | 1.94 |
| HERCULES difference-2 | 0.00 | 0.00 | 0.00 | 1.42 | 4.33 | 3.32 | 21.56 | 0.00 | 3.02 | 1.42 | 3.02 |

In vivo experiments showed successful acquisition and separation of ISTHMUS into PRESS and HERCULES data. Linewidths of tCr and the agreement of linewidth between the first and second scan are shown in Figure 4. One-way ANOVA suggested no significant differences between PRESS and HERCULES data within the same brain region as shown in Figure 4a. Similar linewidths (∼5.5 Hz) were obtained in the CSO, dACC and PCC and highest linewidths were obtained in the thalamus (∼7.5 Hz). Multiple comparisons suggested linewidths in the thalamus were significantly higher (p<0.001) than any of those in the other 3 brain regions. The Bland-Altman plot showed the agreement of linewidth between the two scans. Most of the differences fell within ±1 Hz and the maximum differences were up to approximately ±1.5 Hz as shown in Figure 4b. SNRs were not compared since PRESS and HERCULES data were acquired using different number of transients. Linewidths from phantom and in vivo data suggested high quality data were collected.

**Figure 4a).**
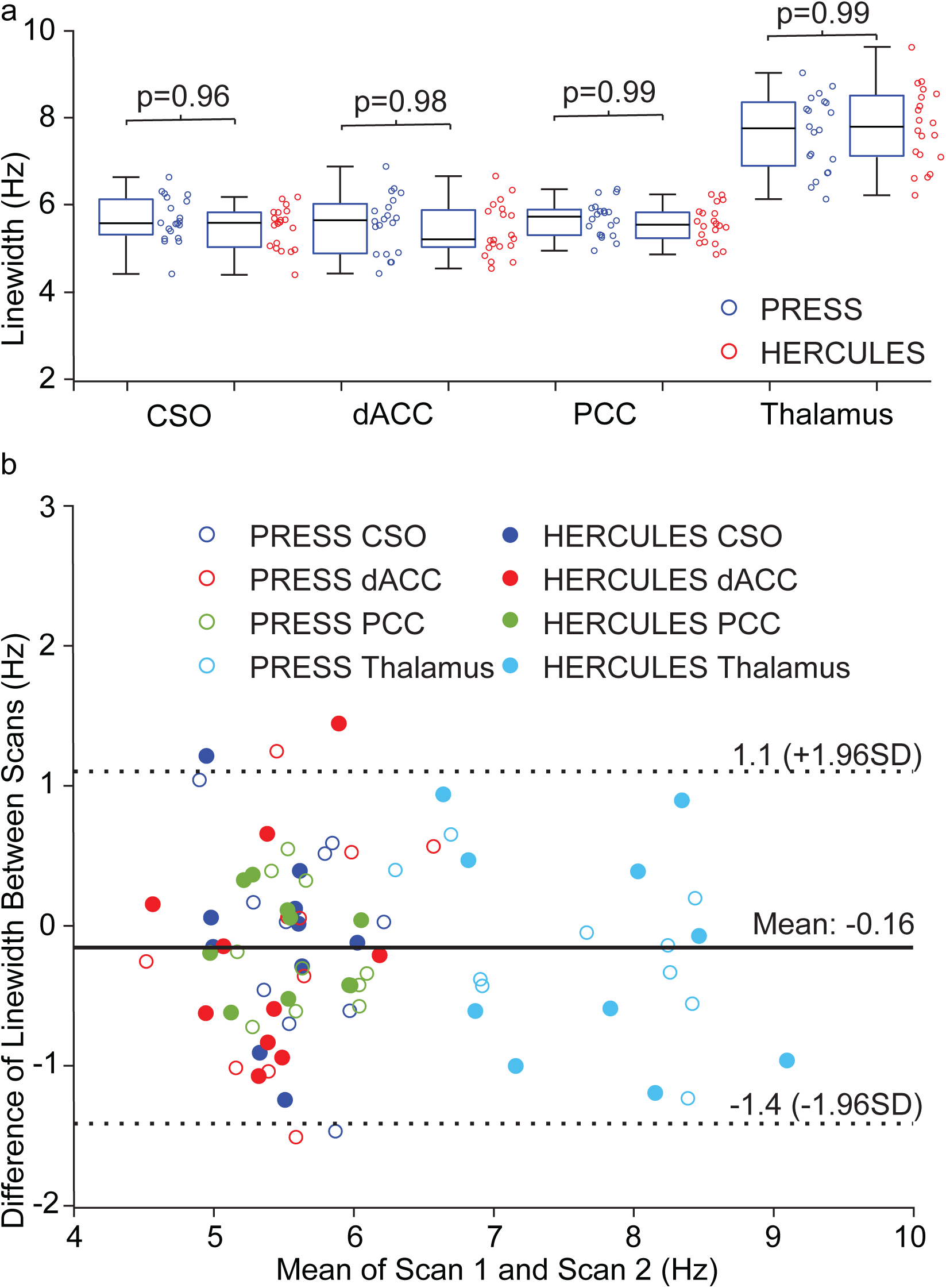
Box plots depicting linewidths of tCr in four brain regions. Multiple comparisons suggested no significant difference between PRESS and HERCULES within the same brain region and linewidths in the thalamus are significantly higher compared to those in the other regions. 4b) shows the Bland–Altman plot for the agreement of linewidth between the first and the second scan.

In vivo ISTHMUS data of PRESS and the three Hadamard combinations (green) of HERCULES overlaid by the linear combination modeling (LCM, yellow) are shown in Figure 5. LCM of the in vivo ISTHMUS spectra are shown in Figure 6. Conventional ‘measurable’ metabolites including tNAA, tCr, tCho and mI were clearly visible in the PRESS data as shown in Figure 6a. Same metabolites were observed in Figure 6b in the HERCULES sum spectrum. GABA was successfully modeled and shown in the GABA-diff spectrum (A+B-C-D) in Figure 6c. Asc, Asp, GSH, Lac, NAAG and PE were modeled in the GSH-diff spectrum (A-B+C-D) in Figure 6d. The LCM of MM and lipid were included but not shown in the figure. Quantitative metabolite concentration measurements and their CVs for the four brain regions are shown in Table 2. In HERCULES, metabolite measurements for NAA, tNAA, tCho, tCr, Gln, Glu, Glx and mI were retrieved from the sum spectrum, GABA from the first diff-spectrum (A+B-C-D) and Asc, Asp, GSH, Lac and NAAG from the second diff-spectrum (A-B+C-D). CVs from the bilateral thalamus were the highest (i.e., most variable) compared with the other three brain regions for most metabolite measurements in PRESS and HERCULES. CVs in the same brain regions between PRESS and HERCULES were relatively similar and comparable for the high concentration metabolites including mI, tNAA, tCho, tCr and Glx. The HERCULES simultaneous modeling approach yielded slightly to moderately lower CVs for Asc, Asp, GABA and Lac compared to PRESS metabolites but was worse for GSH and NAAG potentially due to overlaying of Asc and NAA respectively.

**Figure 5.**
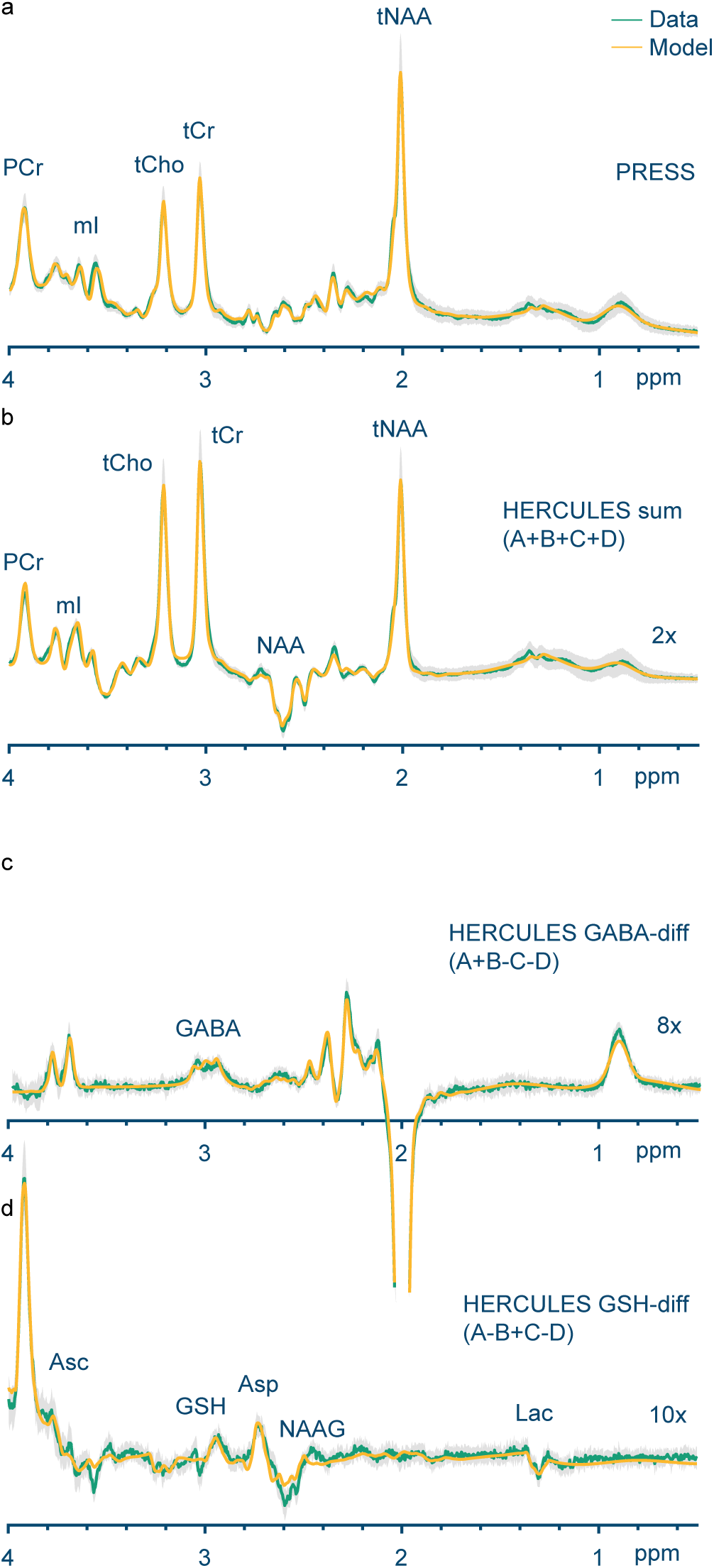
Modeling of the in vivo data from CSO, 5a) showing the short-TE unedited spectrum and the three Hadamard combinations of 5b) sum, 5c) GABA-diff and 5d) GSH-diff. Unedited and edited metabolites are clearly visible.

**Figure 6.**
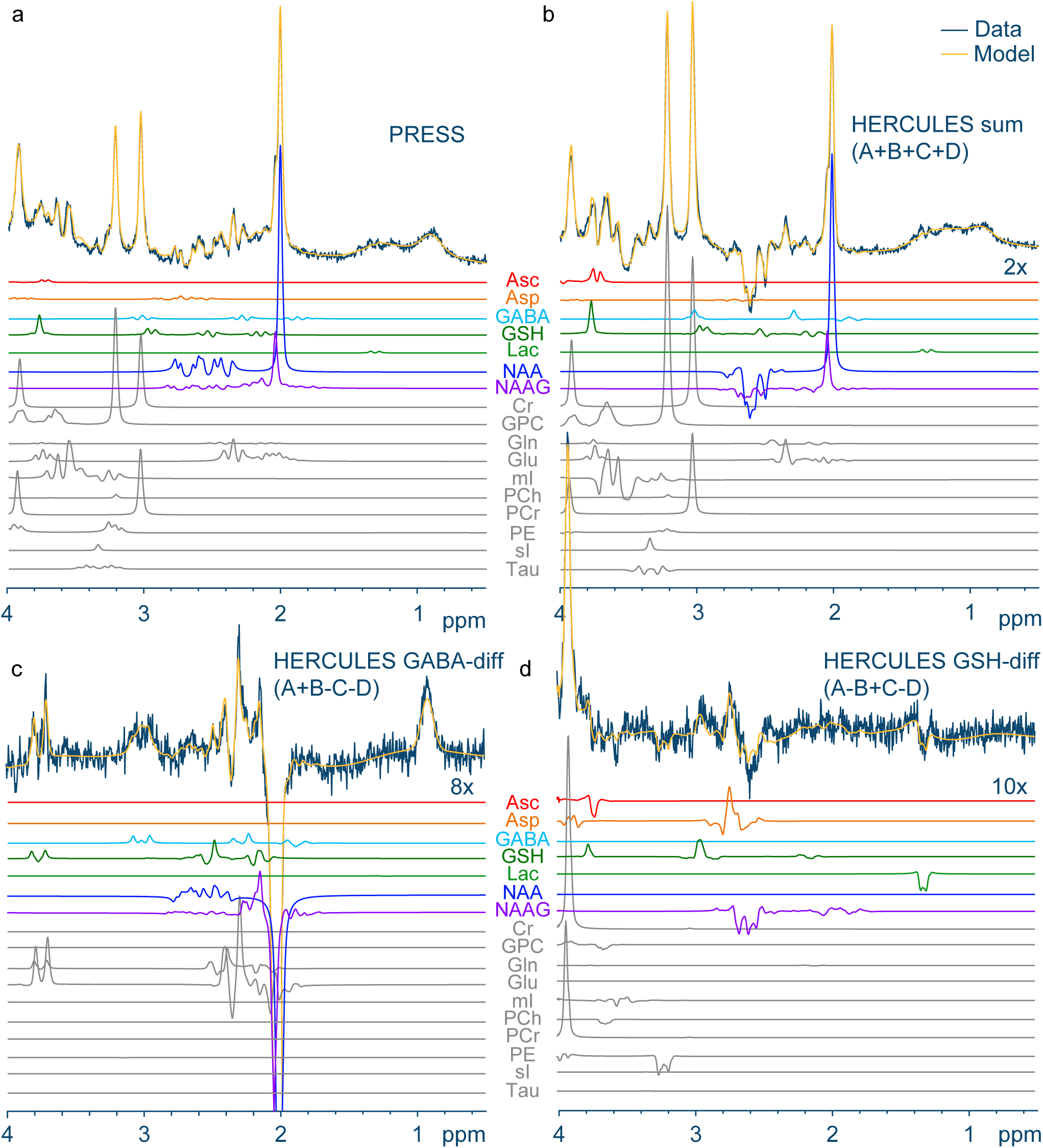
Basis functions of the linear combination modeling of the in vivo short-TE unedited and long-TE edited HERCULES spectra from CSO. Five unedited metabolites (Gln, Glu, mI, tCr, tCho) can be measured in the short-TE (6a) and long-TE sum spectrum (6b) and seven edited metabolites (Asc, Asp, GABA, GSH, Lac, NAA, NAAG) can be measured in the edited spectrum (6c and 6d).

**Table 2.**
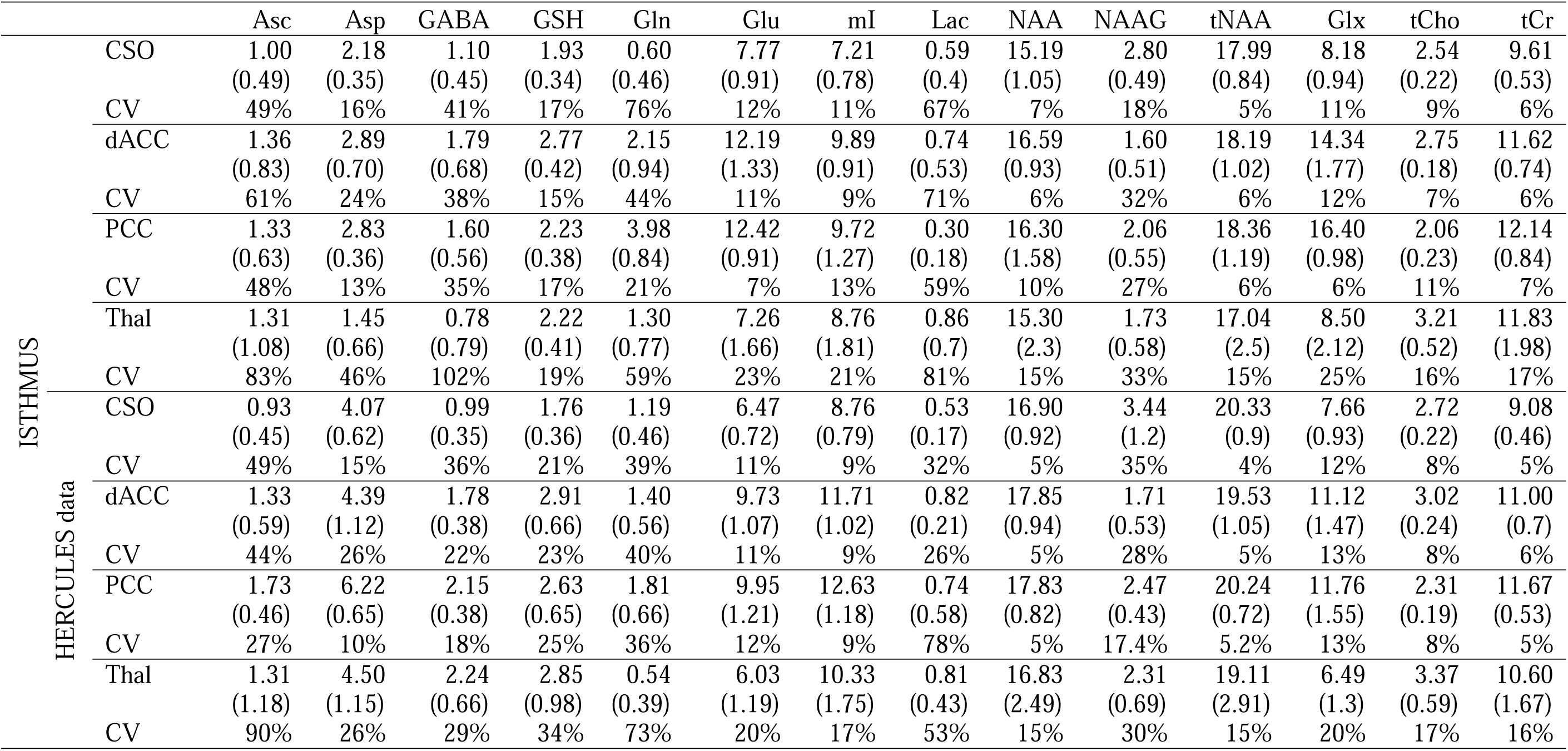
Quantitative metabolite concentration measurements (SD) and coefficient of variation (CV, %) from the in vivo data acquired using ISTHMUS in CSO, dACC, PCC and bilateral thalamus. The data were separated into the PRESS (32 transients) and HERCULES (224 transients) compartments and modeled separately using Osprey. In the HERCULES compartment, measurements for tCho, tCr, Gln, Glu, Glx, mI, NAA and tNAA were acquired from the sum spectrum, GABA from the first difference (A+B-C-D) spectrum and Asc, Asp, GSH, Lac and NAAG from the second difference (A-B+C-D) spectrum. Gln and Glu were quantified separately and combined as Glx. NAA and NAAG were quantified separately and combined as tNAA.

Agreement for metabolite concentration measurements between the first and second scan is shown in Figure 7. Most measurement differences between the two scans were within the 95% confidence interval (1.96 SD) showing good reproducibility of ISTHMUS. Paired t-tests revealed no significant differences in metabolite measurements between the first and the second scans in any of the four regions using the HERCULES data as shown in Supporting Figure S1; one significant difference (p=0.002) was observed in NAAG in the thalamus using PRESS data as shown in Supporting Figure S2.

**Figure 7.**
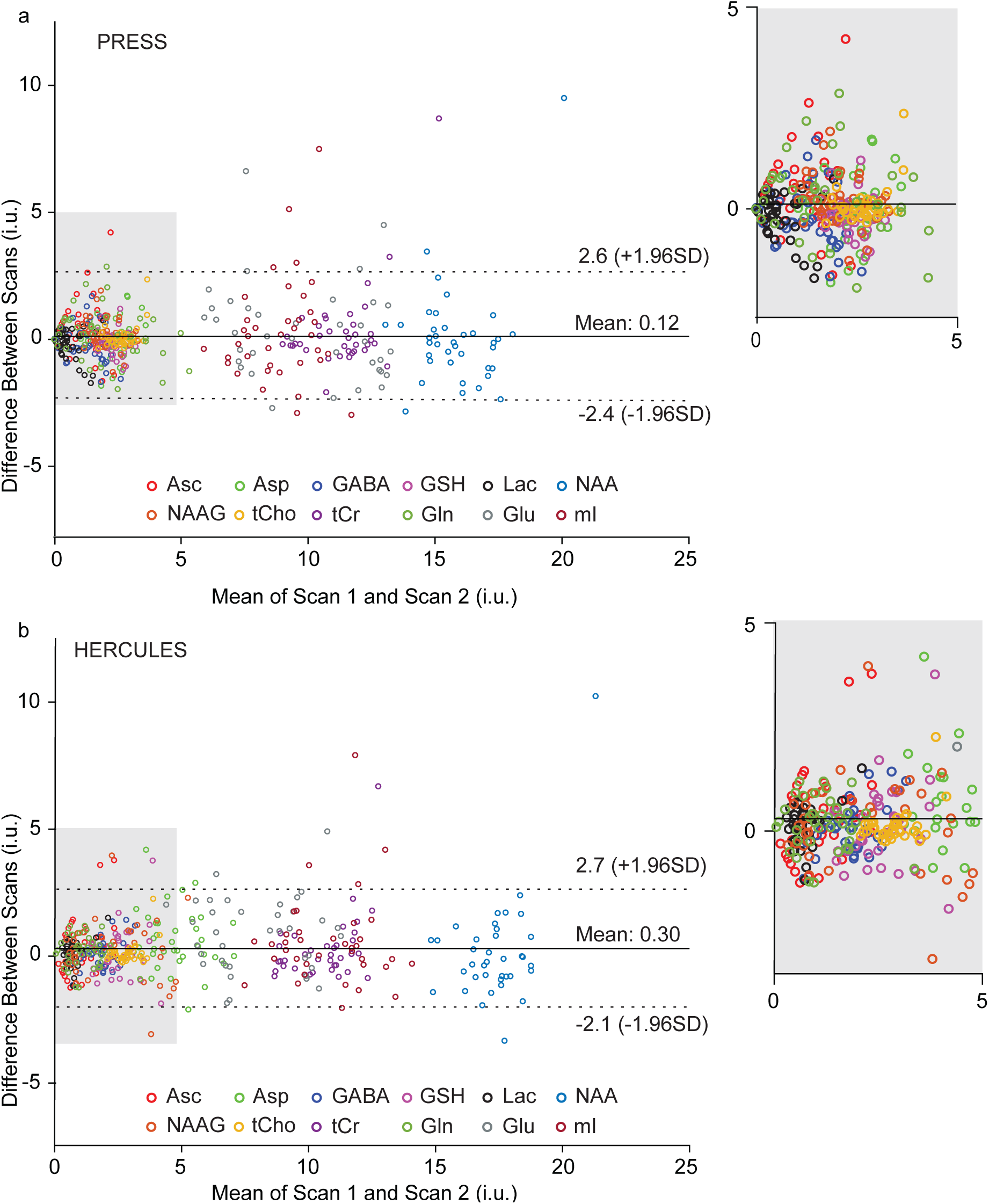
Bland-Altman plot of difference against mean metabolite measurements between the first and the second scan for 7a) PRESS and 7b) HERCULES. Inserts show only those that remain within the gray box on the primary plot to allow for visualization of the lower mean datapoints.

The average water reference signal ratio *R*_integral_ was calculated as 1.87, 1.76, 1.79 and 1.92 (CV: 2.7%, 2.5%, 3.8% and 3.7%) for CSO, dACC, PCC and thalamus, respectively. The average Osprey correction factors *R*_segmentation_ were 1.67, 1.49, 1.46 and 1.57 (CV: 1.3%, 1.7%, 2.3% and 1.8%) for CSO, dACC, PCC and thalamus, respectively based on the default T_2_ values for WM, GM and CSF (Gasparovic et al., 2006; Piechnik et al., 2009). It is noteworthy that the values of *R*_integral_ are larger than those of *R*_segmentation_, and the fractional CVs of are also slightly larger. These values are plotted in Figure 8a, indicating overall correlation in addition to the bias noted.

**Figure 8.**
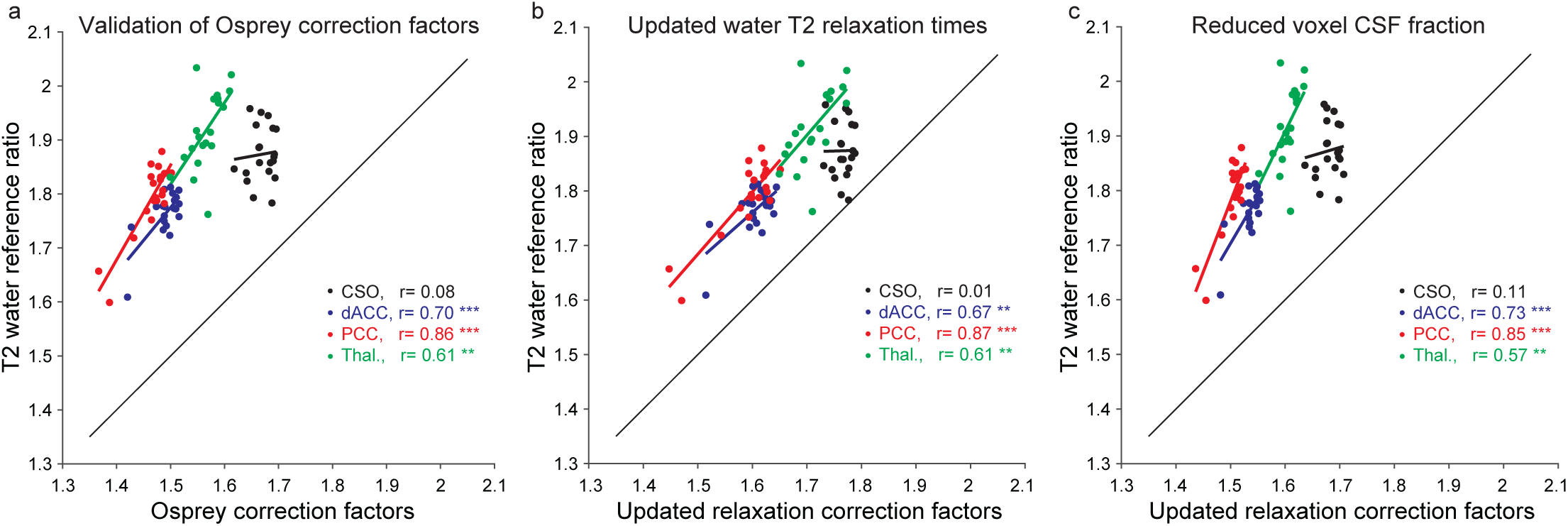
Validation of segmentation-based T_2_ correction, based on literature values. Each panel plots *R*_integral_, the dual-TE water reference ratio against a version of *R*_segmentation_ on the x-axis. Panel a) derives *R*_Segmentation_ from segmented voxel tissue fractions and the Osprey-default literature reference values for water T_2_ in each compartment. Panel b) uses updated literature reference values for water T_2_ to derive *R*_segmentation_. Panel c) uses the Osprey-default literature reference values to derive *R*_segmentation_ but artificially reduces the segmented voxel fractions of CSF by 50%.

The direction of this bias indicates that there is more T_2_ relaxation than the Osprey factors predict between the short- and long-TE water signals. Potential origins of this bias include over-estimation of reference T_2_ values and over-estimation of the voxel CSF fractions. The impact of using shorter T_2_ values for WM and GM resulted in better agreement between *R*_segmentation_ and *R*_integral_, as shown in Figure 8b. Reducing the segmentation fraction of CSF did not greatly impact the bias, as shown in Figure 8c. All measured average water reference signal ratios *R*_integral_ and the average Osprey correction factors *R*_segmentation_ are included in the Supplementary Material 2.

On average, using the dual-TE water reference to estimate a water T_2_ for each subject resulted in a 8.2% reduction in measured metabolite concentrations. The correlation coefficients between concentrations calculated with Osprey reference water T2s and concentrations calculated with single-subject dual-TE estimates were 0.85, 0.89 and 0.96 for tNAA, Cr and Cho, respectively. The correlations between the two concentrations were significantly correlated (p<0.01) to each other as shown in Figure 9.

**Figure 9.**
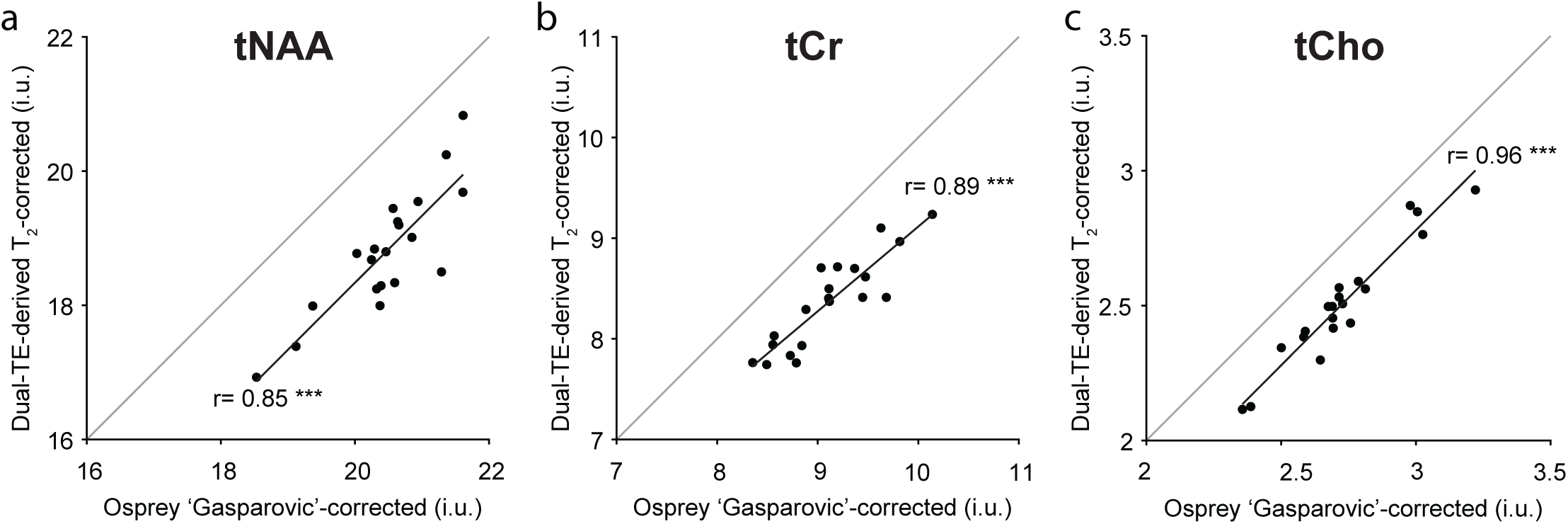
Scatter plot for concentration measurements calculated using dual-TE-derived T_2_ correction against the Osprey default ‘Gasparovic’ correction for a) NAA, b) tCr and c) tCho. The gray diagonal line (y=x) is plotted to visualize bias, and is not a model. Modeled trendlines are shown in black.

## 4 DISCUSSION

ISTHMUS combines short-TE unedited PRESS and long-TE edited HERCULES along with interleaved short- and long-TE water reference scans. Data acquired from CSO, dACC, PCC and bilateral thalamus have been successfully separated into PRESS and HERCULES data and quantified.

The in vivo data were successfully modeled using Osprey. tCho, tCr, tNAA, and mI were clearly visible in the PRESS and HERCULES sum spectrum, while low-concentration metabolites including GABA, GSH, Asc, Asp, PE and Lac were fitted in the HERCULES difference spectra. Similar modeling was observed in the phantom data, with the absence of GSH, Asc and Asp. Quantification of the in vivo data suggested that CVs were low for high-concentration metabolites (i.e. NAA, tCho, tCr, mI) in both PRESS and HERCULES-sum measurements. Bilateral thalamus produced the highest CVs (i.e., most variability) in most metabolites among the four brain regions in PRESS and HERCULES data. Bad shimming, low SNR and potentially larger variations of T_2_ relaxation due to accumulation of iron may contribute to the greater dispersion of the measurements. CVs for edited metabolites such as GABA, GSH, Asc and Lac are in line with a previous literature (Oeltzschner et al., 2019). Asc, Gln and NAAG have relatively high CVs likely due to the overlapping of Cr signal at 3.9 ppm, Glu signal at 3.7 ppm and NAA signal at 2.6 ppm. These overlapping metabolite signals may be ameliorated by improved spectral resolution at higher B_0_ field. Paired t-tests suggested a lack of bias in metabolite measurements between the first and the second scan. No significant differences are observed in HERCULES data and only NAAG measurements from the bilateral thalamus region in PRESS are significantly different between the two scans. This is likely due to the heavy overlap of NAA and NAAG, consistent with the high CVs (>30%) of NAAG in the bilateral thalamus region.

Small linewidths (<6 Hz in CSO, dACC and PCC and <8 Hz in thalamus) and high agreements between the first and second scan (discrepancy < 1.5 Hz) indicate good quality and consistency of data acquisition. The bilateral thalamus yielded significantly higher linewidths (also lower SNR) than any other brain regions in both PRESS and HERCULES data. Multiple factors may contribute to the higher linewidth and the lower SNR including bad shimming, T_2_ relaxation and the potential influence of iron accumulation. Shimming is one of the most essential factors for successful MRS scans. The inhomogeneity of the B_0_ field within the volume of the defined voxel could be deteriorated by the geometry of the biliteral thalamus. The thalamus also has the greatest accumulation of iron besides the basal ganglia and dentate nuclei (McNeill et al., 2012; Peran et al., 2009; Ward et al., 2014). A higher iron concentration leads to shorter T_2_ and T_2_* relaxation and thus broadening of the linewidth and a reduction in the SNR (Wood et al., 2005).

No significant differences (all p>0.96) are observed in linewidth between PRESS and HERCULES within the same brain region as shown in Figure 4a, which show good stability of data quality between the two constituent sequences in ISTHMUS. The Bland-Altman plot also shows good agreement in linewidths between the first and the second scan as shown in Figure 4b. Some outliers are observed in data with lower linewidths (i.e. CSO, dACC and PCC). Since the differences are small (∼1.5 Hz), they are likely contributed by macroscopic B_0_ fluctuations rather than microscopic heterogeneity within the voxel (Juchem and de Graaf, 2017). SNRs have not been compared between PRESS and HERCULES since these scans are acquired with different numbers of transients (32 and 224 respectively).

T_2_ relaxation, which occurs at different rates for metabolite and water signals, must be corrected for in in vivo MRS quantification. Typically, and in community consensus methods (Wilson et al., 2019), this involves using literature reference values for metabolite relaxation rates, and water rates for each major tissue compartment (GM/WM/CSF). However, such assumptions ignore documented T_2_ changes with development, aging and disease, as well as finer-grain spatial heterogeneity across the brain. Errors in assumed relaxation rates propagate strongly into concentration errors especially for in vivo protocols with TE > 30 ms (Gasparovic et al., 2018). The use of reference values for water relaxation times is an enduring limitation of relaxation-corrected MRS concentration calculations, mitigated somewhat by acquiring metabolite and water reference signals that are relaxation-weighted as little as is feasible, i.e. by minimizing TE and maximizing TR. However, reference values are still required, and a renewed effort is merited to improve them. In ISTHMUS, both water and metabolite signals are acquired at two echo times, paving the way for future two-dimensional analyses that are less susceptible to T_2_ errors and changes. It is interesting to note that the dual-TE reference signals are more different than predicted by reference-value T_2_ relaxation corrections in Osprey by ∼10-20%. If transverse relaxation of the water signal during TE is faster than predicted from reference values, this would lead to an underestimation of metabolite levels. It appears that this bias is most likely to come from over-estimation of the tissue T_2_ reference values, since the bias is reduced when more recent, shorter values are used (and not by artificially reducing the voxel CSF fraction). The extent to which reference values are appropriate will vary between studies, and it is possible that greater biases would arise in some studies of older or younger subjects, or in patient cohorts.

Optimized PRESS sequences can achieve echo times as short as 18-20 ms in human brain. The choice of 35 ms here is based upon accessible TE when shortening the edited PRESS sequences used for HERCULES on Philips, GE and Siemens in the HBCD study. In order for the short- and long-TE sequences to be most directly comparable, we chose to maintain the crusher areas across both echo times. One of the motivations of ISTHMUS is to develop an easy, standardized MRS sequence for the multi-site HBCD study. Centers participating in the study include research institutes and local children’s hospitals using scanners from different vendors, in which operators may be less familiar with MRS. Absent integration as ISTHMUS, the collection of edited and unedited MRS data requires 2 scans on Philips and GE (where water reference scans are integrated into each acquisition) and up to 4 scans on Siemens scanners (when water reference scans are additional scans). ISTHMUS helps to integrate all these scans into a single sequence, which ultimately minimizes potential errors from protocol set-up and data export. This allows operators to run the scan in a simple manner once the protocol/exam card has been properly set up. To facilitate and standardize post-processing, ISTHMUS data can be handled by the MRS continuous automated analysis workflow (Zollner et al., 2023a). This end-to-end workflow fully automates data uptake, processing, and quality review, which integrates the most up-to-date file storage and conventions for data acquired using different scanners and vendors. ISTHMUS can be applied in future multi-center studies that use scanners from different vendors other than the HBCD study. Data can be acquired using PRESS and HERCULES to provide a dataset equivalent to ISTHMUS, but care must be taken to ensure appropriate control of voxel placements, receiver gains etc.

Real-time frequency update is implemented into the ISTHMUS scan using the interleaved water reference signal which is particularly essential in a multi-modal MR acquisition protocol, such as the HBCD study, that includes fMRI and DTI sequences prior to the MRS sequence. Both fMRI and DTI sequences require rapid switching of the gradients which generates internal heat to induce frequency drift. A recent study demonstrated that a 30-minute EPI sequence could induce an average frequency drift of more than 3 Hz and the severity varies across scanners with different hardware and cooling systems (Hui et al., 2021). It is essential to have real-time correction to control the quality of the data preventing loss of signal due to spectral broadening and subtraction artifacts from the frequency shift.

The scope of these results is limited to adult data collected from one single scanner of a specific vendor. Data analysis involving multiple platforms would significantly enhance the power of statistics and the conclusion drawn in this study. Ideally, ISTHMUS concentration estimates might be compared to estimates obtained from either or both the standalone short-TE PRESS data and the long-TE HERCULES data. Since each participant underwent two consecutive scans for test-retest purposes, the protocol lasted more than 90 minutes preventing the acquisition of individual PRESS and HERCULES data additionally.

## 5 CONCLUSION

Successful data acquisition using the newly developed ISTHMUS sequence was demonstrated. Quality metrics show consistent data between PRESS and HERCULES. Test-retest analysis shows good reproducibility and high agreement in metabolite measurements. ISTHMUS is less demanding of technologists than its constituent parts separately acquired, hopefully resulting in reduced errors and data loss.

## CRediT authorship contribution statement

**Steve C.N. Hui**: Data curation, Formal analysis, Investigation, Methodology, Software, Validation, Visualization, Writing - original draft, Writing - review & editing. **Saipavitra Murali-Manohar**: Formal analysis, Investigation, Methodology, Software, Validation, Writing - original draft, Writing - review & editing. **Helge J. Zöllner**: Methodology, Software, Validation, Writing - original draft; and Writing - review & editing. **Kathleen E. Hupfeld**: Data curation, Formal analysis, Investigation, Visualization, Writing - original draft, Writing - review & editing. **Christopher W. Davies-Jenkins**: Investigation, Validation, Writing - review & editing. **Aaron T. Gudmundson**: Project administration; Resources, Writing - review & editing. **Yulu Song**: Writing - Data curation review & editing **Vivek Yedavalli**: Project administration; Resources, Writing - review & editing. **Jessica L Wisnowski**: Conceptualization, Investigation, Methodology, Writing - review & editing. **Borjan Gagoski**: Conceptualization, Investigation, Methodology, Writing - review & editing. **Georg Oeltzschner**: Funding acquisition; Investigation, Project administration, Writing - original draft; and Writing - review & editing. **Richard A.E. Edden**: Conceptualization, Funding acquisition, Investigation, Methodology, Project administration, Software, Supervision; Validation; Visualization; Writing - original draft; and Writing - review & editing.

## Declaration of Competing Interest

The authors declare no competing interests.

## Acknowledgement

This work was supported by NIH grants R01 EB016089, R01 EB023963, R01 EB032788, R21 EB033516, R00 AG062230, K00 AG068440, K99 AG080084 and P41 EB031771.

---

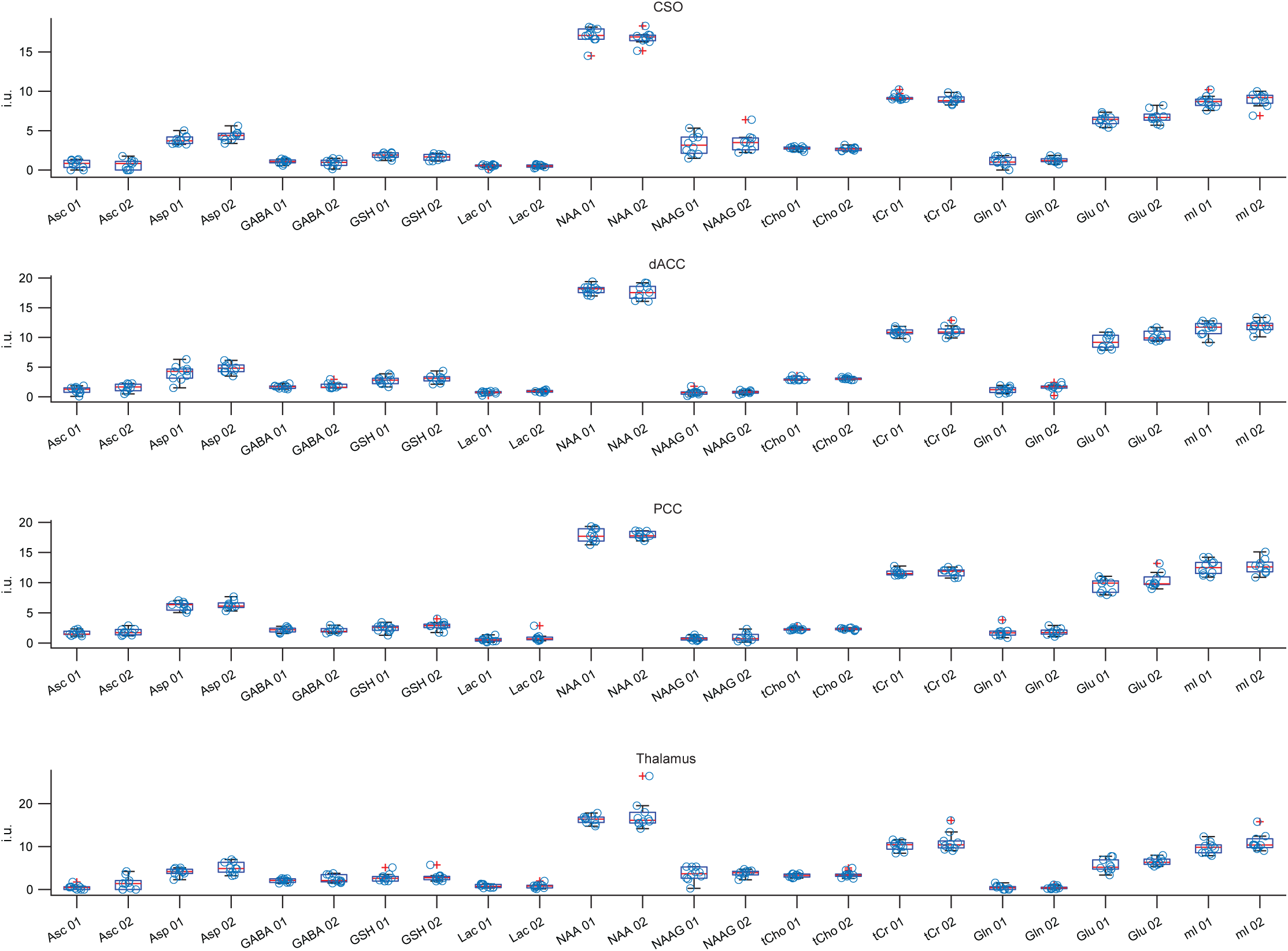
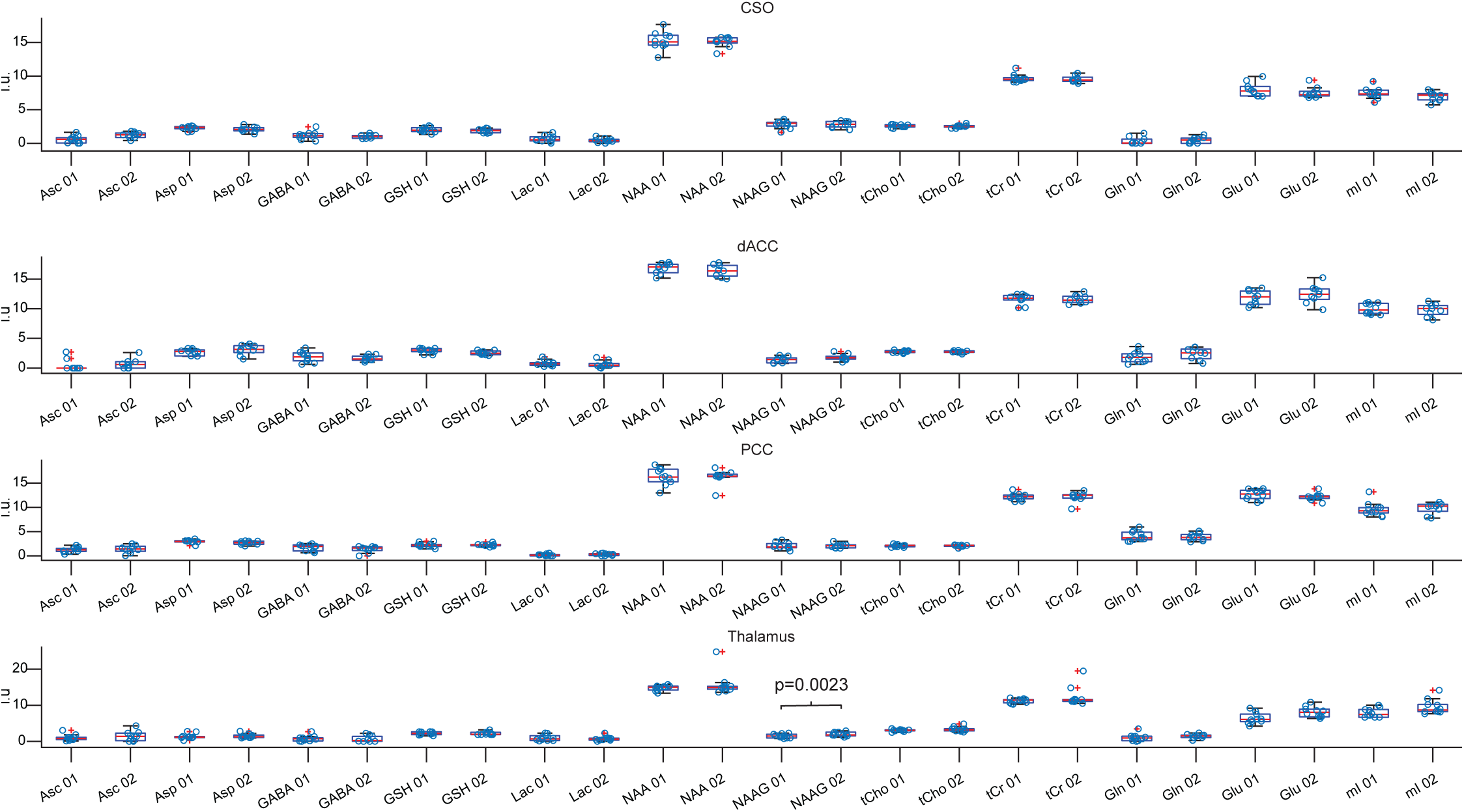

## Notes

### Competing Interest Statement

The authors have declared no competing interest.

### Summary of Updates

Additional analyses for reviews' comments have been made. Figure 8 has been re-plotted and Figure 9 is newly added. Contents of this manuscript have been revised according to reviews' comments.

