## Supplementary data for "Integrated Short-TE and Hadamard-edited Multi-Sequence (ISTHMUS) for Advanced MRS"

**Supplementary data 1:** the minimum reporting standards for in vivo magnetic resonance spectroscopy (MRSinMRS) checklist.

1. Hardware
a. **Field strength [T]:** 3T
b. **Manufacturer:** Philips
c. **Model (software version if available):** R5.71-R2D2
d. **RF coils: nuclei (transmit/receive), number of channels, type, body part:** 32 channel head coil
e. **Additional hardware:** N/A
2. Acquisition
a. **Pulse sequence:** ISTHMUS (i.e. combination of PRESS and HERCULES)
b. **Volume of interest (VOI) locations:** centrum semiovale (CSO), posterior cingulate cortex (PCC), dorsal anterior cingulate cortex (dACC) and bilateral thalamus (Thal.)
c. **Nominal VOI size [cm3, mm3]:** 30×26×26 mm^3^ (CSO/PCC), 30×30×30 mm^3^ (dACC), 25×36×25 mm^3^ (Thal.)
d. **Repetition time (TR), echo time (TE) [ms, s]:** TR: 2000 ms. TE:68/80 ms
e. **Total number of excitations or acquisitions per spectrum:** 32/224 acquisitions (PRESS/HERCULES), 4/4 acquisitions (short-TE/long-TE water)

In time series for kinetic studies

i. Number of averaged spectra (NA) per time point
ii. Averaging method (eg block-wise or moving average)
iii. Total number of spectra (acquired/in time series): 64 increments with 8 averages per increment
f. **Additional sequence parameters (spectral width in Hz, number of spectral points, frequency offsets):** 2000 Hz, 2048 points

If STEAM: mixing time (TM)

If MRSI: 2D or 3D, FOV in all directions, matrix size, acceleration factors, sampling method

g. **Water suppression method:** CHESS
h. **Shimming method, reference peak, and thresholds for “acceptance of shim” chosen:** Automated B_0_ field mapping 2^nd^ order shimming, water linewidth <14 Hz
i. **Triggering or motion correction method (respiratory, peripheral, cardiac triggering, incl. device used and delays):** N/A
3. Data analysis methods and outputs
   a. **Analysis software:** Osprey v2.5.0
   b. **Processing steps deviating from quoted reference or product:** Default segmentation and T2 correction.
   c. **Output measure (eg absolute concentration, institutional units, ratio)**: Absolute concentration.
   d. **Quantification references and assumptions, fitting model assumptions:** N/A
4. Data quality
   a. **Reported variables (SNR, linewidth (with reference peaks)):** SNR and linewidth of the 3-ppm tCr signal.
   b. **Data exclusion criteria:** No subjects excluded.
   c. **Quality measures of postprocessing model fitting (eg CRLB, goodness of fit, SD of residual):** No direct QA measures. Coefficient of variations for test-retest.
   d. **Sample spectrum**: figure 5

**Supplementary data 2**: ratio $R_{\mathrm{integral}}$ (dual-TE water integral) and ratio $R_{\mathrm{segmentation}}$ using default Osprey and shorter T2 values for WM and GM and 50% of the segmented CSF value. $R_{\mathrm{integral}}$for the dual-TE water were plotted against the $R_{\mathrm{segmentation}}$ as shown in Figure 8.

| **Dual-TE water integral** | CSO | dACC | PCC | Thalamus |
| --- | --- | --- | --- | --- |
| Mean | 1.87 | 1.76 | 1.79 | 1.92 |
| SD | 0.05 | 0.04 | 0.07 | 0.07 |
| CV | 2.7% | 2.5% | 3.8% | 3.7% |
| **Default Osprey T2** |  |  |  |  |
| Mean | 1.67 | 1.49 | 1.46 | 1.57 |
| SD | 0.02 | 0.03 | 0.03 | 0.03 |
| CV | 1.3% | 1.7% | 2.3% | 1.8% |
| **Shorter T2** |  |  |  |  |
| Mean | 1.77 | 1.60 | 1.59 | 1.71 |
| SD | 0.02 | 0.03 | 0.05 | 0.04 |
| CV | 1.0% | 2.1% | 3.2% | 2.2% |
| **CSF-50% segmented** |  |  |  |  |
| Mean | 1.68 | 1.54 | 1.50 | 1.60 |
| SD | 0.02 | 0.02 | 0.02 | 0.02 |
| CV | 1.1% | 1.2% | 1.5% | 1.2% |
